# Temperature shapes movement and habitat selection by a heat-sensitive ungulate

**DOI:** 10.1101/790048

**Authors:** Jesse M. Alston, Michael J. Joyce, Jerod A. Merkle, Ron A. Moen

## Abstract

**Context:** Warmer weather caused by climate change poses increasingly serious threats to the persistence of many species, but animals can modify behavior to mitigate at least some of the threats posed by warmer temperatures. Identifying and characterizing how animals modify behavior to avoid the negative consequences of acute heat will be crucial for understanding how animals will respond to warmer temperatures in the future.

**Objectives:** We studied the extent to which moose (*Alces alces*), a species known to be sensitive to heat, mitigates heat on hot summer days via multiple different behaviors: (1) reduced movement, (2) increased visitation to shade, (3) increased visitation to water, or (4) a combination of these behaviors.

**Methods:** We used GPS telemetry and a step-selection function to analyze movement and habitat selection by moose in northeastern Minnesota, USA.

**Results:** Moose reduced movement, used areas of the landscape with more shade, and traveled nearer to mixed forests and bogs during periods of heat. Moose used shade far more than water to ameliorate heat, and the most pronounced changes in behavior occurred between 15°C and 20°C.

**Conclusions:** Research characterizing the behaviors animals use to facilitate thermoregulation will aid conservation of heat-sensitive species in a warming world. The modeling framework presented in this study is a promising method for evaluating the influence of temperature on movement and habitat selection.

## Introduction

Physiological performance peaks within a limited range of body temperatures in which molecular, cellular, and systemic processes operate optimally. Body temperatures outside this range impose functional constraints on these processes, including reductions in growth, reproduction, activity, and immune function (Pörtner and Farrell 2008). Nevertheless, animals routinely operate in environmental conditions that trigger suboptimal body temperatures (Boyles et al. 2011; Sunday et al. 2014). This conundrum underlies two long-standing questions in biological research: (1) How do animals mitigate suboptimal thermal conditions, and (2) how effective are those efforts at mitigation? Rapid and ongoing responses to global climate change by a multitude of animal species (Walther et al. 2002; Parmesan 2006; Hoegh-Guldberg and Bruno 2010) increase the urgency of answering these questions.

Animals can relax the constraints of limited ranges of thermal tolerance by modifying their behavior to reduce heat gain and dissipate heat at high temperatures. Such behavioral thermoregulation has a long history of study in biological research (Cowles and Bogert 1944), but this idea still offers fresh insight today. Animals can restrict movement to produce less metabolic heat (Stelzner 1988; Broders et al. 2012), alter posture to reduce heat gain from insolation or increase surface area to shed heat (Luskick et al. 1978; Bartholomew and Dawson 1979), pant to lose heat via evaporation (Campos and Fedigan 2009; McCann et al. 2013), or visit thermal refugia (spaces that provide refuge from thermal stress caused by extreme temperatures [e.g., burrows, wallows, shade cover]; van Beest et al. 2012; Hovick et al. 2014; Kurylyk et al. 2015), among other behaviors. Identifying exactly which of these strategies animals use to behaviorally thermoregulate, how much these strategies buffer against adverse impacts of hot weather, and what costs animals incur to use these strategies is crucial for understanding their ability to increase ranges of thermal tolerance, which in turn increases our understanding of how animals may adapt (or not) to increasing temperatures in a warmer future.

Recent advances in statistical modeling techniques provide opportunity to study behavioral thermoregulation in new ways. Step-selection functions (hereafter “SSFs”) are an extension of the resource selection function modeling framework that explicitly incorporates spatial and temporal animal movement characteristics to enable examination of fine-scale habitat selection and movement behavior at biologically realistic scales (Forester et al. 2009; Thurfjell et al. 2014; Duchesne et al. 2015; Signer et al. 2019). SSFs have primarily been used to model habitat selection (Thurfjell et al. 2014), but recent theoretical development has demonstrated that they can also be used to explicitly model movement behavior that changes in both space and time in relation to landscape features (Avgar et al. 2016; Prokopenko et al. 2017; Ladle et al. 2019; Signer et al. 2019). By including interaction terms between temperature, habitat covariates, and movement rates within SSFs, the relative importance of temperature-dependent animal behaviors—including both resource selection and movement rates—can be quantified in a single model.

We used the SSF modeling framework to examine behavioral thermoregulation in moose (*Alces alces*), a species known to be sensitive to heat. Moose have undergone substantial population declines across much of their southern range due in part to climate change (Lenarz et al. 2010; van Beest et al. 2012; Dou et al. 2013; Monteith et al. 2015). Moose experience heat stress starting at temperatures as low as 14°C (Renecker and Hudson 1986) or 17°C (McCann et al. 2013) during the summer. Moose prevent heat stress on hot days by using water, shade, and decreased activity to shed heat via conduction and reduced exposure to radiation from the sun (Fig. 1; Belovsky 1981; Dussault et al. 2004; Broders et al. 2012; Street et al. 2015; McCann et al. 2016). At coarse spatial and temporal scales, moose select for thermal cover (e.g., dense canopy in conifer forests) during periods of high temperatures (Schwab and Pitt 1991; Demarchi and Bunnell 1995; van Beest et al. 2012; Melin et al. 2014; Street et al. 2016; but see Lowe et al. 2010). However, earlier studies have not established the relative importance of multiple different heat amelioration strategies (e.g., seeking shade vs. reducing movement vs. visiting water) or identified thresholds at which behavioral thermoregulation alters habitat selection.

To evaluate how moose modify fine-scale habitat selection and movement patterns as temperatures increase, we used an SSF to assess the effects of temperature on movement and resource selection. We examined empirical support for a single model consisting of temperature and interactions with variables likely to be important for moose thermoregulation. This model enabled us to quantify the importance of several ways moose may alter behavior to thermoregulate when it is hot: moose (1) decrease movement rates to decrease metabolic heat production, (2) increase use of shade to decrease heat gain from solar radiation, (3) increase use of water to increase heat loss via conduction, convection, and evaporation, or (4) use some combination of each of these.

## Materials and Methods

### Study area

We conducted our study in northeastern Minnesota, USA. Federal, state, county, and tribal public lands managed for timber harvest and recreation make up >80% of property ownership in the area. The region is a sub-boreal transition zone between northern hardwood forests in the south and Canadian boreal forests in the north (Pastor and Mladenoff 1992). Upland forests are primarily composed of white, red, and jack pine (*Pinus strobus, P. resinosa*, and *P. banksiana*), aspen (*Populus tremuloides*), paper birch (*Betula papyrifera*), and balsam fir (*Abies balsamea*). Black spruce (*Picea mariana*), tamarack (*Larix laricina*), and northern white cedar (*Thuja occidentalis*) dominate wet lowland forests. Mean minimum and maximum temperatures, respectively, are −16.5°C and −5.5°C for the month of January and 12.6°C and 24.0°C for the month of July at the Beaver Bay weather station on the southern edge of our study area (National Oceanic and Atmospheric Administration). Snow cover is typically present from December to April, with mean annual snowfall ranging between 150 – 240 cm (Minnesota Department of Natural Resources).

### Animal Capture and GPS Telemetry

We captured moose by darting them from helicopters (Quicksilver Air, Inc., Fairbanks, Alaska, USA) during the winters of 2011 and 2012. Darts used to sedate moose contained 1.2 ml (4.0 mg ml^-1^) carfentanil citrate (ZooPharm, Laramie, Wyoming, USA) and 1.2 ml (100 mg ml^-1^) xylazine HCl (Midwest Veterinary Supply, Inc., Burnsville, Minnesota, USA), and we used 7.2 ml (50 mg ml^-1^) naltrexone HCl (ZooPharm) and 3 ml (5 mg ml^-1^) yohimbine HCl (Midwest Veterinary Supply) as antagonists (Roffe et al. 2001; Lenarz et al. 2009). We fitted immobilized moose with global positioning system (GPS) collars (Lotek Wireless, Inc., Newmarket, Ontario, Canada). Animal capture and handling protocols met American Society of Mammalogists recommended guidelines (Sikes et al 2011) and were approved by the University of Minnesota Animal Care and Use Committee (Protocol Number: 1309-30915A).

Collars were programmed to record locations every 20 minutes and to drop off moose at the end of expected battery life (2 years). We retained GPS locations with 3-D fixes or 2-D fixes with dilution of precision values ≤ 5 (Lewis et al. 2007) and removed locations that resulted in movement rates > 30 km / hour. Data used in this analysis include only locations between May 1 and September 31—dates that coincide with average daily maximum temperatures above the threshold believed to induce heat stress for moose (Renecker and Hudson 1986). Location and activity data within 2 weeks of death or collar failure were censored from our data, and only full months of data were used in analysis. Our analysis included 153 moose-months from 24 moose. Moose were adults at capture except for one moose that was a yearling (1.8 years old), and 17 of 24 moose were females.

### Model Covariates

Because shade is difficult to directly calculate over large areas at fine scales and varies at any given location on daily and seasonal cycles, we used canopy vegetation density as a proxy for shade. Canopy vegetation density was estimated using airborne lidar data. Lidar is an active, laser-based remote sensing technology that provides detailed information on topography and vegetation structure (Vierling et al. 2008; Davies and Asner 2014). Lidar data were collected over our entire study area during leaf-off conditions in May 2011 as part of the Minnesota Elevation Mapping project (Minnesota Geospatial Information Office). Lidar data were collected from a fixed wing airplane at an altitude of 2,000-2,300 m above ground level using discrete-return laser scanning systems (ALS60, ALS70, or Optech GEMINI). Side overlap was 25% with a scan angle of ± 20°. Nominal point spacing and pulse density varied due to incomplete overlap of adjacent flight-lines. Average nominal pulse density was 1 pulse/m^2^. We calculated height of discrete returns above ground by subtracting ground elevation based on a lidar-derived Digital Elevation Model from the return elevation. Lidar data met the National Standard for Spatial Database Accuracy (Federal Geographic Data Committee 1998) and had a vertical accuracy RMSE of 5.0 cm and a horizontal accuracy of 1.16 m.

We estimated canopy vegetation density as the proportion of all returns that were ≥ 3 m above ground. Canopy vegetation density is typically estimated with LiDAR as the ratio of returns originating from the canopy to the total number of returns (Vierling et al. 2008; Merrick et al. 2013; Davies and Asner 2014). We used a threshold of 3 m to estimate canopy vegetation density because this corresponds to canopy over a moose’s head. Lidar-derived canopy vegetation density estimates were summarized in a 30 × 30 m grid that aligned with 2011 National Land Cover Database (NLCD; Homer et al. 2015) raster data to ensure consistency across data layers in GIS. We used FUSION software (McGaughey 2016) to create the lidar-derived canopy vegetation density raster. For the sake of simplicity, we hereafter refer to lidar-derived canopy vegetation density as “shade”.

Vegetation cover types were determined using the 2011 NLCD (Homer et al. 2015). NLCD is a remotely sensed dataset of 16 land cover classes created from Landsat Thematic Mapper with 30 m spatial resolution. We extracted 5 vegetation cover types that may offer thermal refuge—woody wetland, hereafter called bog; emergent herbaceous wetland, hereafter called marsh; open water; conifer forests; and mixed forests. Each of these cover types offers different amounts of thermal refuge via different mechanisms (Table 1). Each cover type also offers different amounts of forage. Since moose primarily eat the leaves of deciduous shrubs and saplings < 3 m tall during summer (Peek et al. 1976), forage quantity decreases as the amount of shade and proportion of conifers increases. Selection of cover types may be dependent on proximity to other cover types, and GPS error may lead to underestimation of selection of cover types covering small areas (Conner et al. 2003; Martin et al. 2018). We therefore calculated the Euclidean distance of each pixel in our study area from each of our chosen vegetation cover types using ArcMap 10.4 (Esri, Redlands, California, USA). Euclidean distances were 0 when an animal was within the land cover type of interest.

**Table 1.** Variables incorporated in the step-selection function of moose movement and habitat selection in response to changes in temperature and justification for inclusion in the model.

| Name | Variable | Description |
| --- | --- | --- |
| Shade | Canopy Vegetation Density | Proportion of all lidar returns above 3 meters; analogous to canopy vegetation density, a proxy for shade |
| dBog | Distance to Bog | Distance to woody wetlands; included in analyses because bogs have both canopy cover and ground moisture |
| dMarsh | Distance to Marsh | Distance to emergent herbaceous wetlands; included in analyses because moose are often observed in marshes, and water can disperse heat via conduction, convection, and evaporation |
| dWater | Distance to Open Water | Distance to open water; included in analyses because moose are often seen in bodies of water, which can disperse heat via conduction, convection, and evaporation |
| dConifer | Distance to Conifer Forest | Distance to conifer forest; included in analyses because conifer forest contains localized thick canopy cover |
| dMixed | Distance to Mixed Forest | Distance to mixed forest; included in analyses because conifers offer localized thick canopy cover while deciduous trees offer foraging opportunities |
| StepLength | Step Length | Distance between a moose location and the location immediately prior; included in analyses to account for bias in the parametric distribution of step lengths used to characterize available points and to estimate how temperature affects movement rates |
| Temp | Temperature | Temperature at the nearest NOAA weather station at the time of a location; included in analyses to estimate how temperature affects habitat use and movement rates |

Temperature data were obtained from two weather stations within our study area—KBFW in Silver Bay and KCKC in Grand Marais (Fig. A1; MesoWest). These stations are operated by the National Oceanic and Atmospheric Administration according to national standards and report temperatures at 20-minute intervals. Moose locations were individually matched with the nearest weather station (by distance) and nearest temperature recording (by time). Moose locations were on average 33 km from the nearest weather station and 7 minutes from the closest recorded weather observation in time.

### Statistical Analysis

We used a step-selection function (SSF) to model moose resource selection and movement behavior. For our SSF, we selected available points using a parameterized Weibull distribution of step lengths and the observed distribution of turn angles of the animals in our data set. We paired 20 available locations to each used location (i.e., 21 points per stratum). Our final data set contained 311,521 steps taken by 24 moose. We used conditional logistic regression (via the ‘clogit’ function in the ‘survival’ package; Therneau 2019) to fit the SSF containing our variables of interest (extracted at the end of each step; listed in Table 1) and interactions between each variable and ambient temperature. We included step length (i.e., distance between consecutive fixes) both to reduce bias in selection estimates (Forester et al. 2009) and to explicitly model its interaction with another variable of interest (Avgar et al. 2016; Prokopenko et al. 2017; Ladle et al. 2019). Interaction coefficients detail how temperature influences step length and selection of cover types at differing temperatures. Because step lengths vary in a regular pattern over the course of each 24-hour period (Fig. A2), we adjusted step lengths prior to inclusion in the model by subtracting the observed step length from the average step length at each given time of day. Because moose move more during crepuscular periods than at other times of day regardless of temperature (Cederlund 1989; Lowe et al. 2010; Eriksen et al. 2011; Vander Vennen et al. 2016), failure to adjust for crepuscular activity peaks could lead to consistent positive bias in movement rates at low (morning) and intermediate (evening) temperatures. We included one-way interactions between each covariate and temperature (°C). Because temperature was constant within strata, it was considered only as an interaction term. The full final model is listed below:

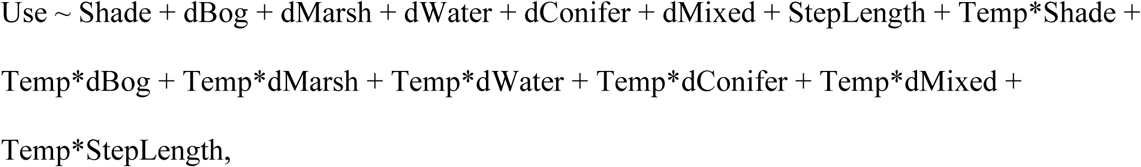

where “*” denotes interactions between variables. We used generalized estimating equations (GEEs) to obtain robust standard errors among animal-days that reduce Type I error caused by pseudoreplication (Fortin et al. 2005; Craiu et al. 2008; Duchesne et al. 2010), and checked that variance inflation factors (VIFs) between main effects were < 3 to ensure multicollinearity was not a problem in our model (O’Brien 2007; Dormann et al. 2013). We calculated VIFs using the ‘vif’ function in the ‘car’ package (Fox and Weisberg 2019). We then conducted k-fold (k=5) cross-validation on our final model and calculated Spearman rank correlation (mean of 50 replications) to evaluate model fit based on the methods of Fortin et al. (2009). Finally, we rarified data to 1-, 2-, and 4-hour intervals to demonstrate the impact of less frequent GPS location data on our ability to detect biologically significant interactions. All analyses were conducted using R statistical software (R Core Team 2018). Effect sizes are reported as relative selection strength (RSS) for one location on the landscape relative to another, given the difference in a variable of interest between the two locations while holding the values of all other variables in the model constant (Avgar et al. 2017).

## Results

### Moose Movement and Resource Selection

We found empirical support for four interaction terms (StepLength*Temp, Shade*Temp, dBog*Temp, dMixedForest*Temp) in our step-selection function (Table 2), indicating that temperature significantly altered movement rate and selection for shade, bog, and mixed forest. We did not detect empirical support for interactions between temperature and distance to marsh, temperature and distance to open water, or temperature and distance to conifer forest. Of these variables with interaction terms whose 95% CIs overlapped zero, only the main effect for distance to conifer forest was significant. Regardless of temperature, moose selected areas further from conifer forest (Relative Selection Strength [RSS] = 1.553; 95% CI = 1.133-2.130). Moose neither selected nor avoided areas near marsh or open water. Habitat use by moose was consistent throughout our study period (i.e., month-to-month changes in distance to vegetation cover types were small; Fig. A3).

**Table 2.**
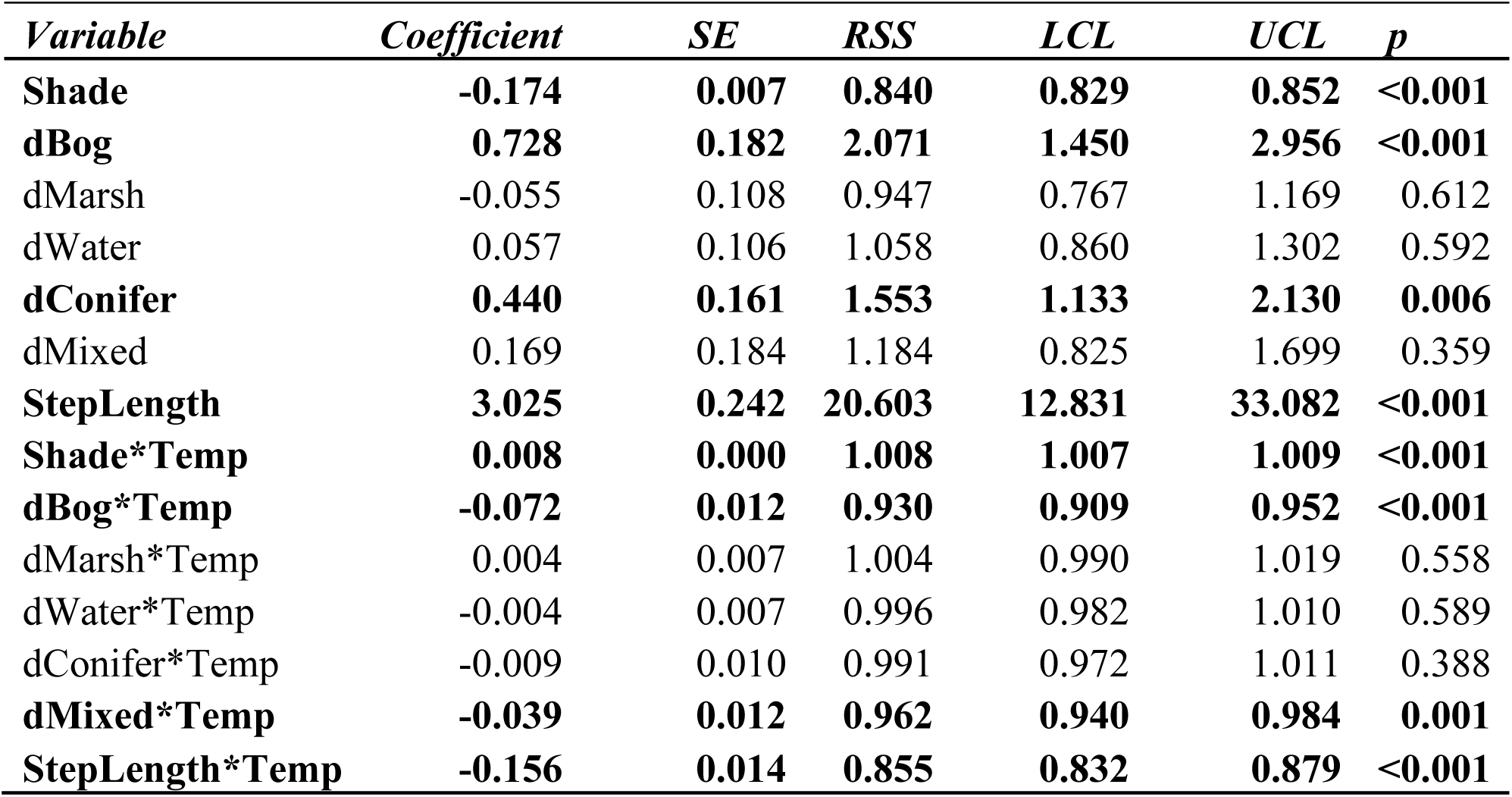
Coefficient estimates, standard errors (SE), relative selection strengths (RSS), 95% confidence limits (lower, LCL; upper, UCL), and p-values for a step-selection function of moose habitat selection and movement in response to changing temperature. Bold lettering denotes variables with confidence limits that do not overlap 1 (the dividing line between selection and avoidance). Variables are described in detail in Table 1.

Moose decreased movement rates at hotter temperatures (Fig. 2A). At each standardized step length > 0 m (i.e., steps that were longer than average for a given time of day), the odds of moose taking a step of that length was higher at 0°C than at 15°C, and higher at 15°C than at 30°C. At 0°C, the odds that moose would move 100 m more than average in 20 minutes were substantially higher (RSS = 1.074; 95% CI = 1.025-1.126) than at 15°C (RSS = 0.849; 95% CI = 0.777-0.928), which were in turn substantially higher than at 30°C (RSS = 0.672; 95% CI = 0.590-0.765). Furthermore, the mean step length at all temperatures above 20°C was below the overall mean step length controlling for time of day.

**Fig. 1.**
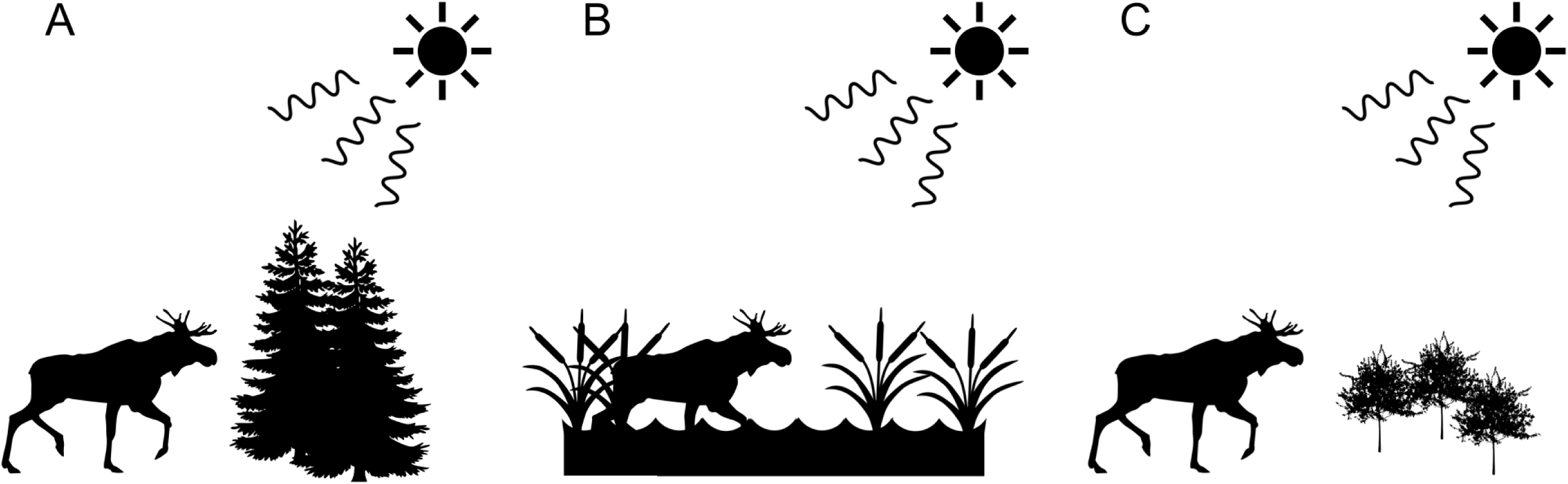
The physical characteristics of the surrounding environment greatly influence the thermal landscape for animals. Fig. 1A represents an environment (conifer forest) where heat gain may be decreased by reducing exposure to radiation, Fig. 1B represents an environment (marsh) where heat loss may be increased by conduction, and Fig. 1C represents an environment (clear cut) that offers neither relief from radiation nor opportunities to disperse heat via conduction. Moose likely face tradeoffs between forage availability and thermal relief.

**Fig. 2.**
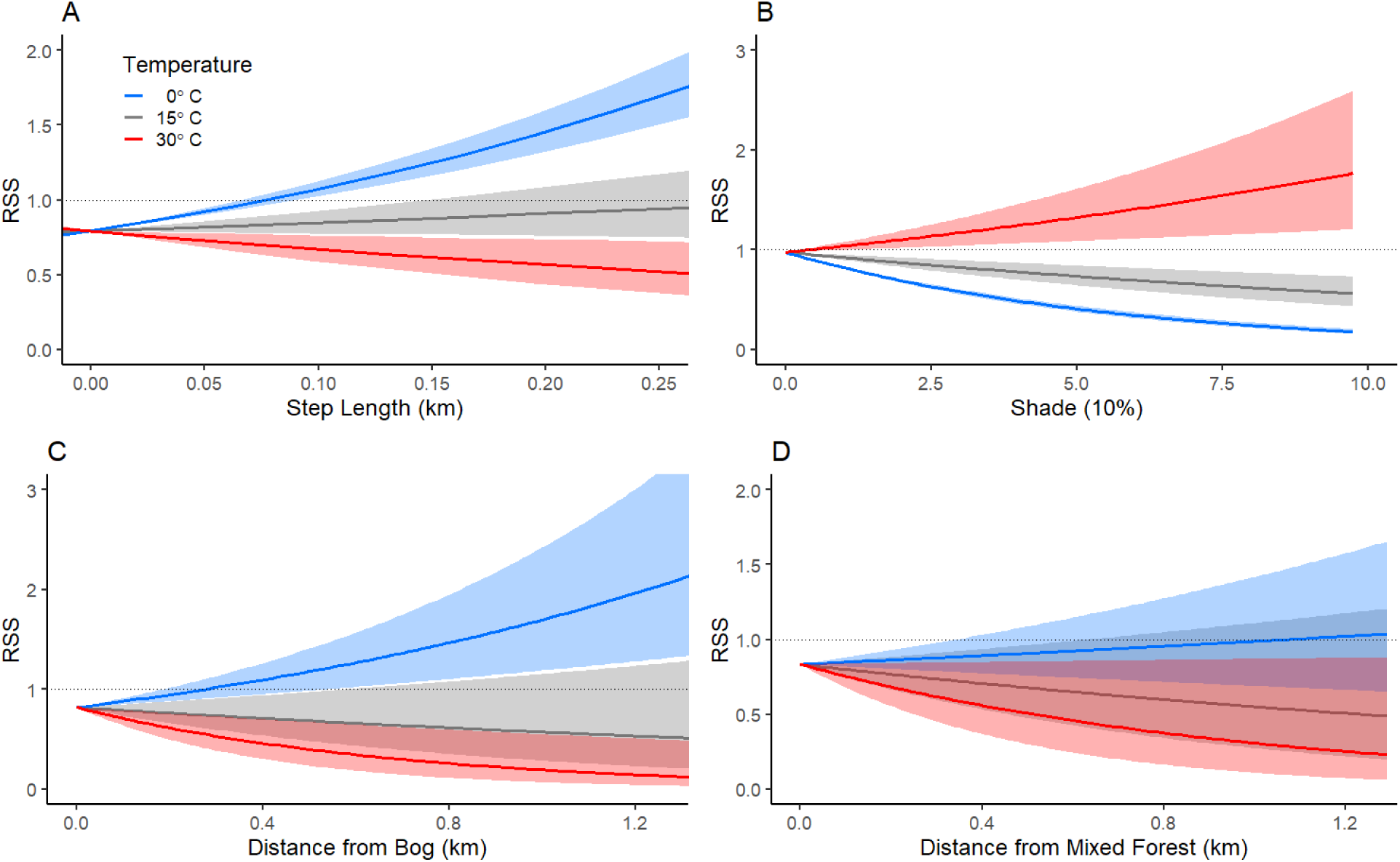
Interaction plots showing relationships for significant interactions between temperature and relative selection strengths (RSS) of variables of interest (A: Step length and temperature, B: Shade and temperature, C: Distance to bog and temperature, D: Distance to mixed forest and temperature). High temperatures decrease the odds of longer step lengths, increase the odds of seeking shade, and increase the odds of traveling in bogs and mixed forest. In some cases (A, B, and C), patterns of behavior at low temperatures reversed into patterns of the opposite behavior at high temperatures (e.g., moose strongly avoid shade at 0°C while strongly selecting for shade at 30°C).

Moose spent more time in shade at hotter temperatures (Fig. 2B). Relative selection strength increased with increasing vegetative cover at 30°C, while it decreased with increasing vegetative cover at 0°C and 15°C, indicating that moose sought shade at high temperatures while avoiding it at lower temperatures. At 0°C, the odds that moose would move into a pixel with 75% vegetative cover were substantially lower (RSS = 0.265; 95% CI = 0.239-0.295) than at 15°C (RSS = 0.640; 95% CI = 0.523-0.782), which were in turn substantially lower than at 30°C (RSS = 1.542; 95% CI = 1.148-2.073).

Despite avoiding bogs at colder temperatures, moose traveled closer to bogs at hotter temperatures (Fig. 2C). The odds that moose were far from bogs was higher at 0°C (RSS = 1.180; 95% CI = 0.988-1.410 at 500 m) than at 15°C (RSS = 0.686; 95% CI = 0.484-0.973 at 500 m), and higher at 15°C than at 30°C (RSS = 0.399; 95% CI = 0.237-0.672 at 500 m).

Moose selected for shorter distances to mixed forest when it was hot than when it was cold (Fig. 2D). The odds that moose were far from mixed forest were higher at 0°C (RSS = 0.909; 95% CI = 0.756-1.088 at 500 m) than at 15°C (RSS = 0.679; 95% CI = 0.478-0.963 at 500 m) or 30°C (RSS = 0.507; 95% CI = 0.302-0.852 at 500 m), though only the difference between 0°C and 30°C was statistically significant.

### Model validation

K-fold cross-validation of our model demonstrated that our model was substantially better than random at predicting where moose moved—the mean Spearman rank correlation coefficient was 0.47 for observed steps.

### Effects of temporal scale on interactions

The interactions we found between temperature and step length, shade, distance to bog, and distance to mixed forest diminish substantially if GPS locations are rarified so that locations occur at longer intervals. When 20-min interval GPS data are rarified to 1-hr, 2-hr, and 4-hr intervals and used to fit the same SSF, interactions become progressively less biologically meaningful (Fig. A4). As the length of time intervals between locations increases, differences across temperatures for step length, shade, and distance to bog become minimal. Differences across temperatures for distance to mixed forest shrink, but more gradually.

## Discussion

In this paper, we developed a modeling framework to test multiple competing (but not mutually exclusive) hypotheses on behavioral responses by animals to heat. We used this framework to model behavioral responses by moose, an ungulate known to be sensitive to heat (Renecker and Hudson 1986; McCann et al. 2013; Melin et al. 2014). Moose altered both movement and habitat selection to behaviorally thermoregulate during hot periods. Moose reduced movement and moved nearer to or stayed within shade, bogs, and mixed forest at high heat, even while avoiding shade and bogs at cooler temperatures (Fig. 2). This pattern links previous findings of separate studies. First, moose prefer to forage in areas with low canopy cover, likely because canopy cover is generally inversely related to forage availability (Lone et al. 2014). Second, moose prefer to use bed sites under dense forest canopy in wet lowland forests during the day (McCann et al. 2016), where moose have access to less forage but more protection against heat gain from solar radiation and more capacity to lose heat to wet ground via conduction. Moose therefore face a steep tradeoff during periods of heat—areas that are good for foraging may not be good for avoiding heat. Selection for shade and shorter step lengths as temperatures increase indicates that moose forego foraging in favor of bedding down under shade as temperatures increase. Earlier studies have documented moose shifting activity to cooler evenings and nights on hot days (Dussault et al. 2004; Montgomery et al. 2019), which is consistent with this trade-off.

The vegetation cover types used more by moose during warm weather further indicate that moose face a tradeoff between foraging and thermoregulation during periods of heat. In general, moose are more likely to find greater quantities of forage in cover types that do not provide thermal cover, while cover types that provide thermal cover are less likely to provide forage. For example, upland mixed forest has some available forage, but forage availability is highest in this cover type in young forests with little canopy cover. Similarly, bogs in Minnesota are largely populated by black spruce, tamarack, and alder, all of which can provide thermal cover but are very rarely eaten by moose (Peek et al. 1976). Birch (*Betula* spp.), willow (*Salix* spp.), and red-osier dogwood (*Cornus sericea*) are eaten by moose and occasionally grow in bogs in Minnesota, but rarely at densities high enough to compensate for unpalatable species dominating the canopy layer.

Further studies would be helpful for demonstrating how common trade-offs between thermoregulation and foraging are among ungulates. Reductions in activity during periods of heat are widespread among ungulates, having been documented in a diverse array of ungulates that include moose, mule deer (*Odocoileus hemionus*; Sargeant et al. 1994), white-tailed deer (*O. virginianus*; Wolff et al. 2020), bighorn sheep (*Ovis canadensis*; Alderman et al. 1989), Alpine ibex (*Capra ibex*; Aublet et al. 2009; Mason et al. 2017), Alpine chamois (*Rupicapra rupricapra*; Mason et al. 2014), common eland (*Taurotragus oryx*; Shrestha et al. 2014), black wildebeest (*Connochaetes gnou*; Vrahimis and Kok 1993), blue wildebeest (*Connochaetes taurinus*; Shrestha et al. 2014), impala (*Aepycerus melampus*; Shrestha et al. 2014), and greater kudu (*Tragelaphus strepsiceros*; Owen-Smith 1998). Nevertheless, the relative importance of reducing activity is not often compared directly to other strategies used by ungulates to reduce heat stress, and reductions in activity may not always result in reduced foraging opportunity, which depends on landscape structure (i.e., distance between foraging sites and bed sites with adequate thermal cover).

Studies on the contexts in which tradeoffs between foraging and thermoregulation become particularly acute will also be important in a warming future. Environment and nutritional condition may play a role in shaping such tradeoffs. For example, North American elk (*Cervus canadensis*) prioritize reducing thermoregulatory costs over forage quality in low-elevation desert populations but not in high-elevation mountain populations, and individuals with low fat reserves prioritize reducing thermoregulatory costs over forage quality most strongly (Long et al. 2014). Body size may also play a role in modulating tradeoffs between foraging and thermoregulation. Adult male Alpine ibex, which are substantially larger than females, reduce time spent foraging more than females when it is hot (Aublet et al. 2009). Common eland and blue wildebeest reduce afternoon activity all year round, but smaller impala reduce afternoon activity only during the summer (Shrestha et al. 2014). The effects of environmental variation, nutritional condition, or body size on thermoregulatory behavior could be answered in larger data sets using our modeling framework by building SSFs for each individual and testing for statistical effects of a variable of interest (e.g., fat reserves, body size) on the RSS of a variable of interest (e.g., step length at a given high temperature).

Determining the relative importance of features on the landscape for mitigating heat stress will also be important in a warming future. In our study, moose used shade far more than water to ameliorate heat during hot weather. Moose are commonly observed in bodies of water, and anecdotal evidence suggests that moose use water to shed heat (Schwab and Pitt 1991; Demarchi and Bunnell 1995). Our analysis, however, indicates that moose do not often use open water and marsh to mitigate heat stress; they prefer to seek shadier vegetation cover types. Nevertheless, they do increase use of woody bogs—where both shade and some water are usually available—as temperatures increase. This is consistent with a previous study (McCann et al. 2016) that found that moose prefer bed sites with both canopy cover and relatively high soil moisture. Other ungulates may use different features of the landscape to mitigate heat stress. For example, mountain goats (*Oreamnos americanus*) may move closer to persistent snow cover during hot weather rather than seeking shade (Sarmento et al. 2019). Step-selection functions that include interactions with temperature offer a simple way to test for the relative importance of a wide variety of different landscape features for thermoregulation.

Frequent GPS locations enabled us to detect responses to heat by moose and may explain why previous attempts to characterize moose movement patterns failed to reveal a strong relationship between temperature and movement rates (Dussault et al. 2004; Montgomery et al. 2019). Moose spend about half of the day foraging during the summer, with foraging bouts interspersed by periods of rumination at bed sites. Periods of rest and rumination are typically distinct and occur at regular intervals of roughly 2 hours (Renecker and Hudson 1989; Moen et al. 1996). As the interval between GPS locations increases, the chance that both ambulatory foraging bouts and stationary ruminating bouts are aggregated into a single GPS fix increases, which homogenizes step lengths (Moen et al. 1996). Frequent GPS locations reduce the probability of this happening. Indeed, when our location data was rarified to 1-, 2-, and 4-hour intervals, effect sizes of interactions between temperature and movement rates were progressively reduced (Fig. A4). Because many species have idiosyncratic movement behaviors, movement studies may require intervals between GPS locations within a specific range to best answer research questions concerning animal movement. This is an important consideration for researchers planning studies of animal movement. Researchers should carefully consider the frequency of GPS locations before deploying GPS collars and recognize that GPS data that is too sparse may result in Type II error.

Our analysis can directly inform management and conservation actions for wildlife. Many moose populations at the southern edge of their distribution (including our study area) have undergone substantial declines in the past decade (Lenarz et al. 2010; van Beest et al. 2012; Dou et al. 2013; Monteith et al. 2015). Our results suggest that in a warmer future proximity to shade will strongly influence habitat suitability for moose in areas with abundant forage due to timber harvest and other anthropogenic disturbance. Moose will likely benefit from management action to explicitly promote maintenance of shade near large patches of forage. Because moose prioritize shade over forage when it is hot, moose will likely not feed in large forest openings on hot days (though moose may feed in unshaded forest openings at night; Dussault et al. 2004). Moose will likely spend more time foraging in forest openings with patches of canopy cover than in large homogeneous forest openings. For example, most of the forage in large clearcuts may be inaccessible to moose during hot periods unless the clearcuts contain “reserve patches”, or interior islands or fingers of forest extending into the clearcut. These reserve patches will likely be most helpful for moose if they consist of bog or mixed forest.

Some measure of fitness (or a proxy for fitness) would make it possible to directly link behavioral strategies to a population-level response to large-scale drivers like climate change. Although behavioral thermoregulation mitigates some metabolic costs of hot weather, forgoing foraging to avoid high body temperatures may result in decreased fat reserves, lower fitness, and ultimately in population declines compared to a cooler baseline scenario where moose do not need to behaviorally thermoregulate. Although we did not link behavior to fitness in this study, identifying and quantifying patterns of behavior allows researchers to explicitly test for effects on fitness in subsequent studies. For example, development of conceptual and modeling frameworks to identify and quantify “green wave surfing” behavior (by which animals migrate along paths of rapidly greening forage) in migratory ungulates (Bischof et al. 2012; Merkle et al. 2016; Aikens et al. 2017) allowed researchers to later quantify a connection between green wave surfing and fitness (Middleton et al. 2018). Our study could be used as a foundation for further analyses along these lines, or to parameterize mechanistic models of moose energetic balances under various climate scenarios, land management strategies, or disturbance regimes to project the outcomes of conservation actions taken to benefit moose.

Our analysis demonstrates that advances in animal tracking, remote sensing, and modelling techniques allow us to study responses by free-ranging animals to weather in the field at finer scales than previously possible. SSFs in particular are a valuable tool to answer questions concerning behavioral responses by free-ranging animals to changes in weather in a relatively simple and intuitive way. Because SSFs estimate selection conditionally at each GPS location, each location or step can be connected with a distinct time and spatial location, enabling inference on how animals change movement and habitat selection in space and time in response to specific stimuli. SSFs have been used to characterize animal movements in relation to landscape features, such as grizzly bear (*Ursus arctos*) response to human activity (Ladle et al. 2019) and North American elk, African wild dog (*Lycaon pictus*), and wolverine (*Gulo gulo*) response to roads (Abrahms et al. 2016; Prokopenko et al. 2017; Scrafford et al. 2018). Likewise, SSFs that incorporate interactions between temperature and other variables of interest can characterize changes in movement behavior and habitat use in response to differences in temperature.

In conclusion, moose altered both movement and habitat selection to behaviorally thermoregulate during hot periods by reducing movement rates and increasing use of shaded vegetation cover types that they avoided at cooler temperatures. Moose did not regularly use water sources that lack canopy cover to shed heat. Moose face a tradeoff between forage and thermal cover at high temperatures and forego foraging in favor of seeking thermal cover. Behavior changed at thresholds near (though somewhat above) previously documented heat stress thresholds (Renecker and Hudson 1986; McCann et al. 2013): step lengths decreased at temperatures above 20°C, and selection patterns for shade reversed above 15°C. Future research characterizing strategies for behavioral thermoregulation and consequences of those strategies for fitness will aid conservation in a warming world, for both moose and other heat-sensitive species.

## Acknowledgements

Many thanks to B. Olson, W. Chen, and others who helped collect field data, as well as J. Rick, B. Brito, F. Molina, B. Maitland, S. Esmaeili, and J. Goheen for providing helpful comments on early drafts of this manuscript. This study was funded by Minnesota’s Environmental and Natural Resources Trust Fund, the University of Minnesota-Duluth Integrated Biosciences graduate program, and the University of Wyoming Department of Zoology and Physiology.

## Author Contributions

JA and RM conceived and designed the study; RM collected the data; JA, MJ, and JM analyzed the data; JA led the writing of the manuscript. All authors contributed critically to manuscript drafts and gave final approval for publication.

## Data Availability

Data and R scripts are archived on *Zenodo* at: http://doi.org/10.5281/zenodo.3872407.

## Appendix A: Supplementary Data

**Fig. A1.**
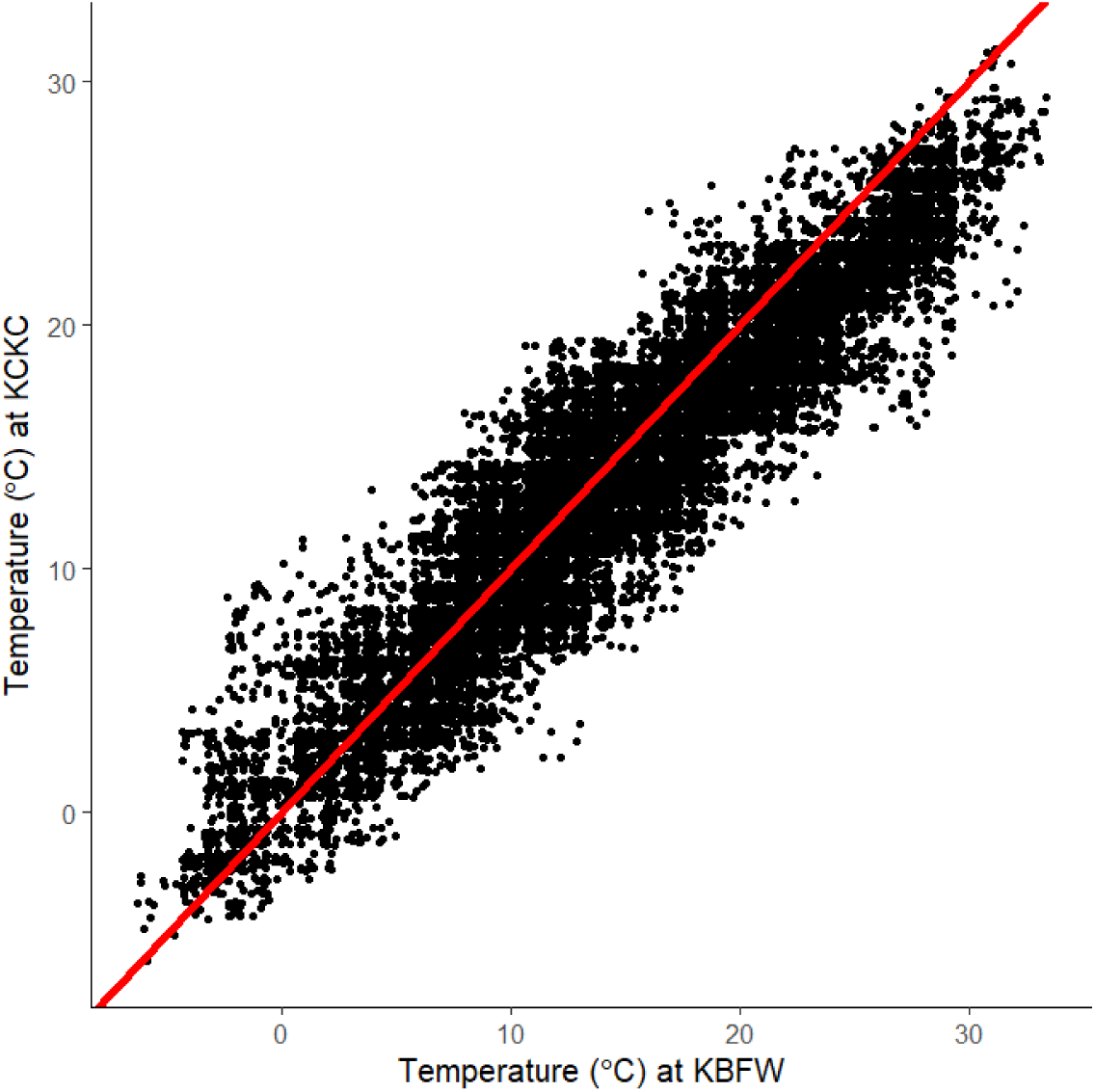
Comparison of temperatures at the two weather stations used in this study (KCKC in Grand Marais and KBFW in Silver Bay). The red line indicates a 1:1 relationship. Temperatures at KCKC followed the regression line 1.74 + 0.821*KBFW, where “KBFW” indicates the temperature at the KBFW station. R^2^ = 0.881 for the regression equation. Temperatures were thus slightly warmer at KCKC at very low temperatures (less than ∼2°C), but usually slightly cooler (e.g., when it was 30°C at KBFW, the expected temperature at KCKC was 26.4°C). Variation in temperature across our study area was thus much smaller than temperature across the day or summer.

**Fig. A2.**
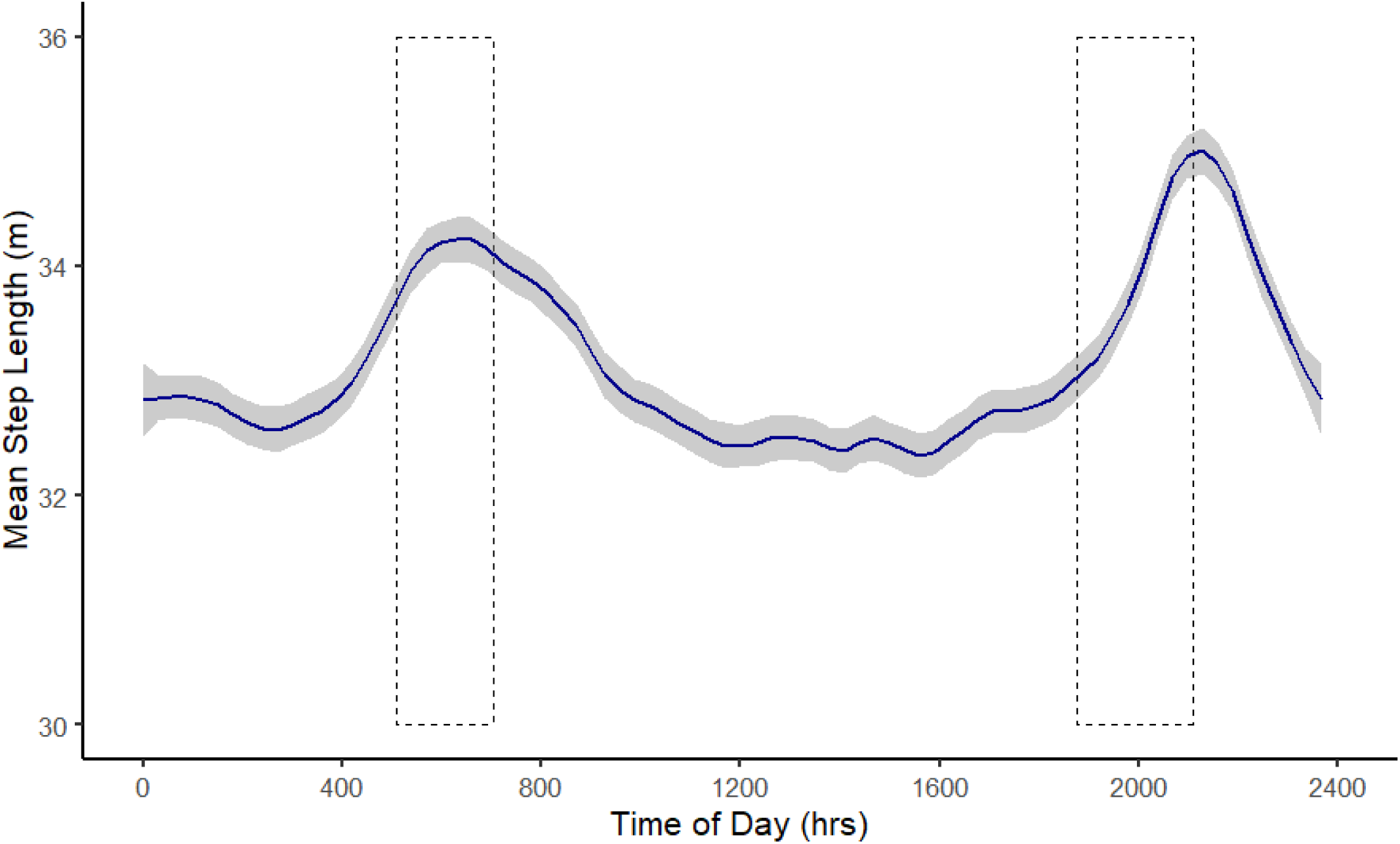
Lowess regression of mean step length across times of day (20 min. increments). The gray ribbon represents the 95% confidence interval for the regression line. Moose movement rates varied slightly but consistently over the course of the day, with movement rates peaking during crepuscular periods. The area within the dotted rectangles represents the range of civil sunrise and sunset at the centroid of our study area during our study period (determined using the NOAA Solar Calculator tool [https://www.esrl.noaa.gov/gmd/grad/solcalc/]).

**Fig. A3.**
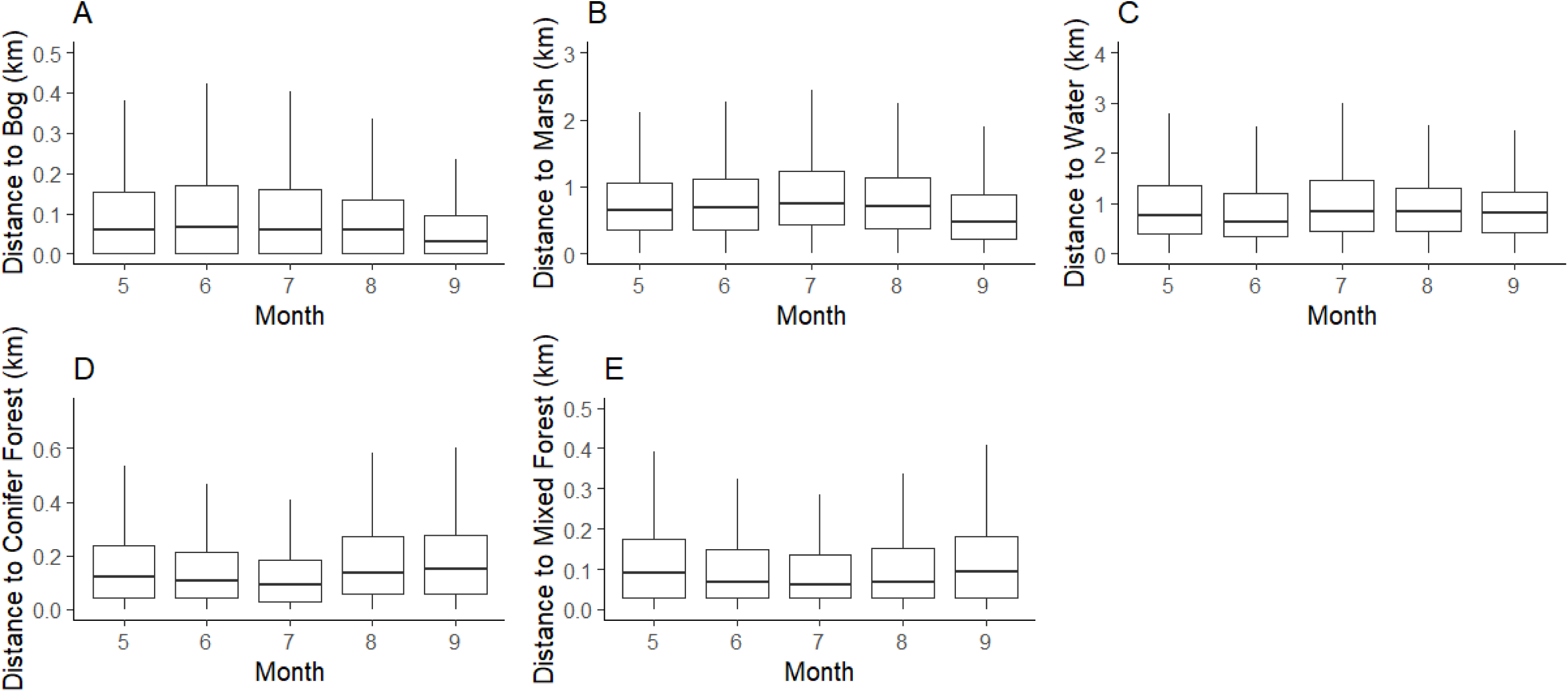
Boxplots showing monthly changes in distance to cover types of interest (A: Distance to bog, B: Distance to marsh, C: Distance to water, D: Distance to conifer forest, E: Distance to mixed forest). Month-to-month differences in habitat use are small, indicating that patterns observed from our SSF reflect habitat selection throughout our study period and are not influenced by one-time phenological events occurring during our study period (e.g., parturition movements by females during May, or the emergence of aquatic plants during June).

**Fig. A4.**
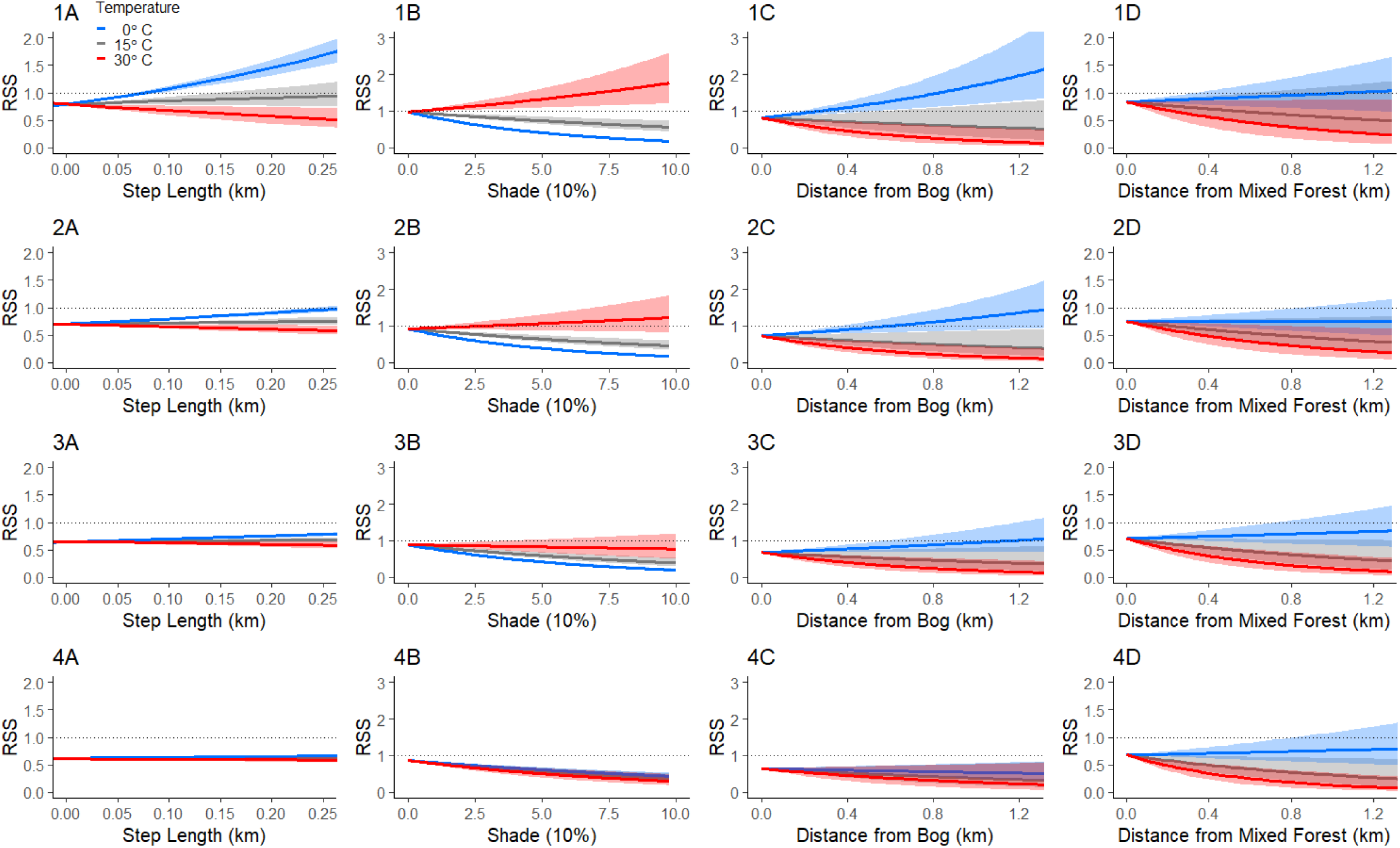
Interaction plots showing relationships for significant interactions between temperature and relative selection strengths (RSS) of variables of interest (A: Step length and temperature, B: Shade and temperature, C: Distance to bog and temperature, D: Distance to mixed forest and temperature) at progressively longer intervals between GPS locations (1: 20-minute, 2: 1-hour, 3: 2-hour, and 4: 4- hour). Patterns in the selection strength of interactions progressively weaken as the interval between GPS locations increases, in part explaining why other studies have not found consistent effects of temperature on moose movement.

